# Phenotypic convergence in the brain: distinct transcription factors regulate common terminal neuronal characters

**DOI:** 10.1101/243113

**Authors:** Nikos Konstantinides, Katarina Kapuralin, Chaimaa Fadil, Luendreo Barboza, Rahul Satija, Claude Desplan

**Affiliations:** Department of Biology, New York University, New York, NY 10003, USA; New York University Abu Dhabi, Saadiyat Island, Abu Dhabi, UAE; New York Genome Center, New York, NY 10013, USA

## Abstract

Transcription factors regulate the molecular, morphological, and physiological characters of neurons and generate their impressive cell type diversity. To gain insight into general principles that govern how transcription factors regulate cell type diversity, we used large-scale single-cell mRNA sequencing to characterize the extensive cellular diversity in the *Drosophila* optic lobes. We sequenced 55,000 single optic lobe neurons and glia and assigned them to 52 clusters of transcriptionally distinct single cells. We validated the clustering and annotated many of the clusters using RNA sequencing of characterized FACS-sorted single cell types, as well as marker genes specific to given clusters. To identify transcription factors responsible for inducing specific terminal differentiation features, we used machine-learning to generate a ‘random forest’ model. The predictive power of the model was confirmed by showing that two transcription factors expressed specifically in cholinergic (*apterous*) and glutamatergic (*traffic-jam*) neurons are necessary for the expression of *ChAT* and *VGlut* in many, but not all, cholinergic or glutamatergic neurons, respectively. We used a transcriptome-wide approach to show that the same terminal characters, including but not restricted to neurotransmitter identity, can be regulated by different transcription factors in different cell types, arguing for extensive phenotypic convergence. Our data provide a deep understanding of the developmental and functional specification of a complex brain structure.

## Introduction

Different cellular characteristics define unique cell types. Before the molecular revolution, cell types were distinguished by their morphology (Cajal, 1915; Fischbach and Dittrich, 1989; Morante and Desplan, 2008), and whenever possible, by their function and physiology (Kepecs and Fishell, 2014). The advent of RNA sequencing technologies has allowed us to revisit cell type classification, identify new cell types and corroborate pre-existing ones (Macosko et al., 2015; Shekhar et al., 2016). Transcription factors are activated by other transcription factors or by signaling pathways and activate downstream effector genes, thereby shaping these cell types. Therefore, transcription factors represent the core of cell type identity by controlling structural (Santiago and Bashaw, 2014; Sivachenko et al., 2013), molecular (Hobert, 2016; Lee and Pfaff, 2001), and physiological (Juarez-Morales et al., 2016; Kratsios et al., 2015) properties of cells. Thus, a fundamental question in developmental biology is: How do transcription factors generate cell types with specific characteristics?

In *Drosophila,* temporal, spatial, and morphology transcription factors are expressed at different developmental stages (Bayraktar and Doe, 2013; Bertet et al., 2014; Brody and Odenwald, 2000; Enriquez et al., 2015; Erclik et al., 2017; Isshiki et al., 2001; Li et al., 2013) to activate terminal selectors that regulate the genes that define the morphology and physiology of adult neurons (Hobert, 2008, 2011). Combinations of terminal selectors define distinct neuronal cell types and regulate terminal effector gene batteries, maintaining the identity of a neuron long after its specification (Arlotta and Hobert, 2015; Doitsidou et al., 2013; Hobert, 2008, 2011; Park et al., 2016; Skromne et al., 2007; Zembrzycki et al., 2015). This has led to the proposal of the existence of a ‘core regulatory complex’ of transcription factors, which *(i)* establishes cell types and *(ii)* can lead to the evolution of new cell types, when their expression is altered (Arendt et al., 2016). This core regulatory complex shapes neuronal diversity in two timescales: first, by implementing their identity over the course of development and, second, by diversifying cell types over millions of years of evolution. Therefore, the transcription factor underpinnings of neuronal morphology, physiology, and molecular identity are at the intersection of the evolutionary and developmental history of a neuron.

The identification of transcription factors in neurons has been driven by tool availability. Screens have relied on existing of antibodies (Erclik et al., 2017; Li et al., 2013) or reporters for specific transcription factors (Hobert et al., 2016), which restricted the search to already known factors. Although numerous terminal selectors and other transcription factors that play essential roles in generating diverse neurons have been identified, the advance of single cell sequencing technologies (Wang and Navin, 2015; Ziegenhain et al., 2017) offers the opportunity to ask the same question in an unbiased way for every neuronal type: Which transcription factors define cell type identity and how do they regulate their downstream effectors?

Only recently have we started to comprehend molecularly the huge diversity of mammalian neurons (Macosko et al., 2015; Mo et al., 2015; Poulin et al., 2016; Wichterle et al., 2013). In parallel, systematic work in the *Drosophila* optic lobe, which consists of ~70,000 cells, has provided an in-depth and nearly exhaustive description of more than 100 cell types (Cajal, 1915; Fischbach and Dittrich, 1989; Morante and Desplan, 2008; Nern et al., 2015). Therefore, the *Drosophila* optic lobe represents an ideal system to seek a mechanistic understanding of the way transcription factors regulate neuronal identity. The optic lobe is formed by four neuropils (lamina, medulla, lobula, and lobula plate) that are organized in ~750 columns, which correspond to the ~750 ommatidia (unit eyes) of the retina. Unicolumnar neurons occupy each column, while multicolumnar neurons cover a larger receptive field, receiving information from more than one ommatidium. Each unicolumnar cell type is therefore represented by ~750 cells of identical morphology and, probably, very similar molecular identity, while identical multicolumnar neurons are represented fewer times.

We currently have a near complete understanding of how different neuronal types are specified from a pool of neural stem cells (neuroblasts) in the developing optic lobe. The intersection of temporal and spatial transcription factors in the neuroblasts, as well as Notch-driven binary cell fate decisions of the intermediate progenitors (ganglion mother cells) and apoptotic cell death are responsible for most, if not all, of the observed neuronal diversity (Bayraktar and Doe, 2013; Bertet et al., 2014; Erclik et al., 2017; Isshiki et al., 2001; Li et al., 2013; Pinto-Teixeira et al., 2016; Rossi et al., 2017; Suzuki et al., 2013). However, it still not understood how a neuron differentiates once specified, *i.e.* how it acquires the characters that endow its identity.

Although a fair number of Gal4 drivers that label a single cell type in the optic lobe are now available (Jenett et al., 2012; Morante and Desplan, 2008; Nern et al., 2015; Pfeiffer et al., 2008), most of the neuronal types in the optic lobes remain molecularly inaccessible. Single cell RNA sequencing techniques, such as Drop-seq, have revolutionized our access to different cell types by offering an unbiased way of sampling single cells from the tissue of interest and determining their transcriptomes (Karaiskos et al., 2017; Macosko et al., 2015). Moreover, coupling Drop-seq to available genetic tools in *Drosophila* can help us establish causal relationships between transcription factors and their downstream effectors.

We combined Drop-seq of single neurons and RNA sequencing of FACS-sorted single cell types to obtain the transcriptome of all neuronal and glial cells in the adult optic lobes. We used the FACS-sorted cell type-specific sequencing data to identify many biologically meaningful clusters in the single cell clustering. We annotated additional clusters using Gal4 lines for marker genes specific to each cluster. A ‘random forest’ machine learning model was trained from the data and was used to predict the expression of terminal genes based on the transcription factor profile of each cell, allowing us to establish causal relationships between transcription factors and the genes that participate in the generation and release of neurotransmitters. Notably, we find that each cell type uses distinct combinations of transcription factors to regulate the expression of effector genes. Altogether our study provides a detailed understanding of the developmental and functional specification of nearly all neurons in a complex brain structure, and sets the groundwork to address the evolutionary and developmental origin of this diversity.

## Results

### Drop-seq of single neurons and glia in the Drosophila adult optic lobe

To obtain an unbiased characterization of all cell types and their transcriptomes, we used Drop-seq (Macosko et al., 2015) to sequence a large number of single cells from the *Drosophila* adult optic lobes. Drop-seq relies on a microfluidic device to isolate single cells in aqueous drops where each cell is lysed and its mRNAs are marked with a specific barcode (Figure 1A). The samples are then pooled and sequenced together. Drop-seq has a number of advantages: *(i)* it grants unbiased access to the transcriptome of every cell in the tissue of interest without the need for any labeling, *(ii)* it can provide the transcriptome of yet unidentified cell types, and *(iii)* it is cost-effective.

**Figure 1:**
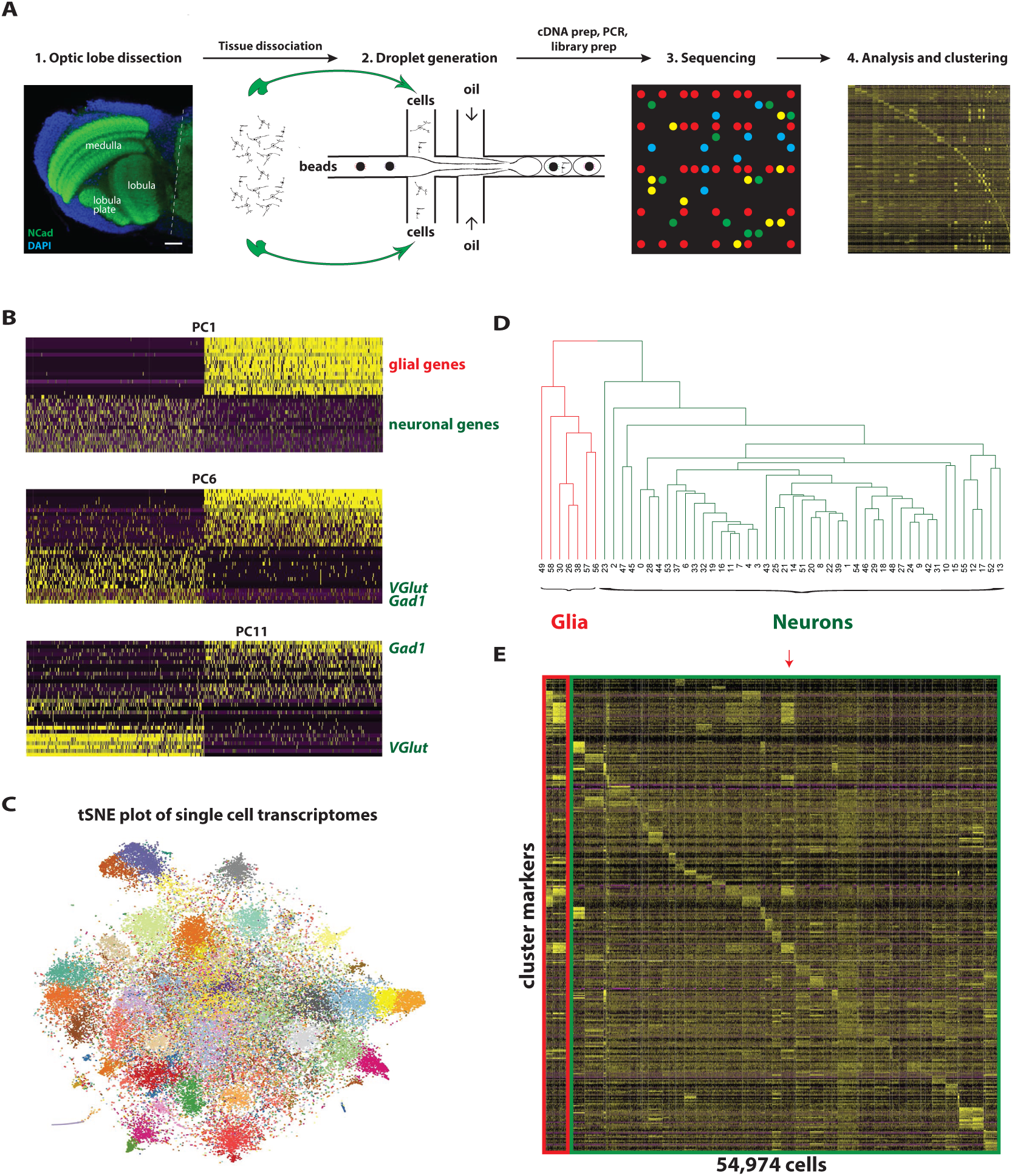
Overview of the Drop-seq experimental procedure, analysis, and clustering. (A) We dissected the optic lobes of the *Drosophila* central nervous system and dissociated them into single cells. We excluded the laminas, and included the medulla, lobula, and lobula plate cells, whose cell bodies can be seen by the DAPI staining in the cortex and rim of the three neuropils (visualized using an antibody against *NCad).* The single cells were then fed into the microfluidic device, alongside the beads (which were in lysis buffer) and the oil, in a setup that resulted in the generation of aqueous droplets in an oil background. Each droplet may be empty, carrying a bead and a single cell, or carrying one of the two. After lysis, transcript annealing, droplet breakage, cDNA preparation and PCR amplification, we sequenced the pooled single cell transcriptomes, analyzed the results using the Seurat package in R and clustered the single cells in 52 clusters. Scale bar, 20um. (B) We performed PCA to reduce the dimensions of the data for further analysis. Genes (rows) and cells (columns) are ordered according to their PCA scores and the 500 most extreme cells and 30 most extreme genes on both sides of the distribution are shown in the heatmap. The first PCs (as indicated here by PC1) were responsible for the separation of neurons from glia, as indicated by the positive contribution of glial genes (such as *nrv2, Inx2, alrm,* and *ogre)* in PC1 and the opposite one for genes enriched in neurons *(VGlut* and nicotinic acetylcholine receptors). Later PCs divide the neurons based on their neurotransmitter identity, as can be seen for PC6 (glutamatergic and GABAergic neurons are separated from the rest, mainly cholinergic ones) and PC11 (glutamatergic and GABAergic neurons are separated from each other). (C) The tSNE plot of all single cells included in our analysis shows the separation of different clusters. We used a k-nearest neighbor algorithm to call 61 clusters, which are shown in different colors on the tSNE plot. (D) Transcription factor-based hierarchical clustering of the Drop-seq cluster transcriptomes. Clusters are numbered from 0 to 58. The first split of the tree represents the separation of 7 glial clusters (red) from 45 neuronal ones (blue), as expected from the PCA. Numbers at the bottom of the tree indicate clusters. (E) The expression of 401 selected Drop-seq cluster markers (rows) is shown in all Drop-seq single cells (columns). Clusters are separated by white lines and are arranged according to the tree in Figure 1D. Glial clusters are highlighted in red, while neuronal clusters are in blue. Interestingly, a single neuronal cluster that expresses elav but not repo (cluster 14 - red arrow) shares many common markers with glia (see Discussion).

*Drosophila* neurons are markedly smaller than in vertebrates. Their diameter is ~3um, while it is ~10um in mice (Shekhar et al., 2016), which results in an approximately 30-fold volume difference. For this reason, cDNA recovery after a Drop-seq run and reverse transcription was too low to allow library preparation with current protocols (Figure S1A) (Macosko et al., 2015). We adjusted the protocol (see STAR Methods) and increased cDNA yield >10-fold (Figure S1A).

We sequenced the transcriptome of ~120,000 adult optic lobe cells and used Seurat (Satija et al., 2015) for the downstream analysis. Using simulated single cell data (see STAR Methods and Figure S1B), the minimum gene-per-cell cutoff was defined at 200 genes. After eliminating cells enriched for mitochondrial gene expression (indicative of stressed cells), our dataset comprised 57,601 cells averaging 538 reads per cell, ranging from 269 to 13,720. 1,303 variable genes were used to perform principal component analysis (PCA). Glial genes contributed significantly to the first, second and third PCs, indicating a clear separation of glia and neurons (Figure 1B). The next PCs separated neurons from each other based on their terminal identity (expression of genes involved in neurotransmitter synthesis and delivery, neurotransmitter receptors, cell adhesion molecules etc). For example, PC6 separated glutamatergic and GABAergic neurons from the other neurons, while PC11 split glutamatergic from GABAergic neurons (Figure 1B). We identified 26 significant PCs following the same jackstraw-inspired procedure used in Macosko et al., (2015). These PCs were used as independent “metagenes” to cluster the single cells. Using a k-nearest neighbor algorithm in PCA space, we identified 61 transcriptionally distinct clusters (STAR Methods, Figure S1C). To visualize the clusters, we reduced the dimensionality of our data below 26 dimensions using t-SNE (t-Distributed Stochastic Neighbor Embedding), a non-linear dimensionality reduction technique (Figure 1C). To evaluate the robustness of the clustering and avoid over-clustering (*i.e.* a single cell type split in two clusters), we followed two different approaches. First, we assessed all terminal nodes of a hierarchical clustering tree by training ‘random forest’ classifiers on each of these nodes and calculating the specificity of the classifier (Figure S1C). Second, for each node for which the error of the classifier was higher than 15%, we looked for differentially expressed genes and assessed whether they justify the split (STAR Methods and Figure S1C-L). This allowed us to eliminate clusters 5 and 36 (enriched in heat-shock proteins indicating stressed cells) and to merge clusters 23 and 40, 29 and 35, 11 and 50, 20 and 41, and 18 and 34. We also eliminated the two smallest clusters (59 and 60), as they only contained 35 and 20 single cells, respectively. We ended up with 52 clusters for further analyses.

To visualize the relationship between the clusters, we performed hierarchical clustering using transcription factor expression as a metric (Figure 1D). As expected from the PCA, the first split differentiated glia and neurons (see below and Figure 3 for details). We also identified specific markers for each of the Drop-seq clusters using Receiver Operating Characteristic (ROC) curves. The expression of the markers for each single cell and cluster is presented in the heatmap of Figure 1E. Most of the markers were not unique to a specific cluster, which was expected as different neuronal types share gene batteries for performing specific functions (Achim and Arendt, 2014; Hartwell et al., 1999). Nonetheless, most of the clusters could be distinguished by the expression of a combination of genes.

In summary, this unbiased technique identified 52 reliable clusters comprising a total of 54,974 cells that represent a large proportion of the optic lobe cellular diversity.

### RNA sequencing of single neuronal types from adult optic lobes

In parallel to Drop-seq, we obtained the bulk transcriptome of 17 neuronal cell types from the adult optic lobes. Gal4 and LexA lines were crossed to flies bearing either a *UAS-RedStinger* or a *LexA-2xhrGFP.nls* transgene to uniquely mark single neuronal cell types. This allowed us to FACS sort the fluorescent cells and sequence their transcriptome in three biological replicates (Figure S2A-B).

We thus sequenced the transcriptome of these 17 cell types that are a representative fraction of optic lobe neurons. They comprise unicolumnar and multicolumnar neurons, local and projection neurons, and cholinergic, GABAergic, and glutamatergic neurons (Figure S2A). PCA verified that the three biological replicates clustered tightly together (Figure S2B).

### Alignment of FACS-sorted and Drop-sequenced neuronal transcriptomes and annotation of Drop-seq clusters

A concern that arises from clustering cells using Drop-seq data is whether these clusters actually represent neuronal and glial cell types, or if the clustering is based on other parameters that could introduce biases. Differentially tuning the parameters of the k-nearest neighbor algorithm can over-or under-cluster the single cells. To address this concern, we turned to the FACS purified RNA-sequenced cell type data. If the single cells were clustered based on their neuronal identity, each of the characterized neuronal types should correspond to one cluster.

To compare the two datasets, we used genes differentially expressed amongst the Drop-seq clusters. A caveat for this comparison is that the two datasets were obtained using vastly different methods. To account for these differences, we simulated single cell data originating from the FACS-sorted cell type-specific transcriptomes. Each “simulated single cell” had a number of total reads generated randomly from a normal distribution centered on the average of the real single cell data with an equal standard deviation. Each read was randomly assigned to a gene with probabilities decided by the cell type-specific transcriptomes.

We then plotted the expression of the cluster-specific markers that were recovered previously on a heatmap and compared the “simulated single cell” heatmap from each FACS-sorted cell type (Figure S2C) to the real single cell Drop-seq clusters (Figure 1E). Figures 2A and S2D show that 15 out of the 17 genetically labeled neuronal types mapped to one unique cluster, verifying the accuracy of our single cell clustering. The remaining 2 cell types (Tm5c and Dm12 neurons) mapped to two or more clusters (Figure S2E). Pearson correlations between the simulated single cell transcriptomes of FACS-sorted cells and the transcriptomes of Drop-seq clusters supported the visual matching (Figure 2B and S2E). In general, the correlation between cell types and clusters was higher for cells types that are represented by more neurons in each brain (unicolumnar neurons), as opposed to less abundant cell types (multicolumnar neurons) and cell types that are heterogeneous (e.g. Tm5ab consists of 2 cell types).

**Figure 2:**
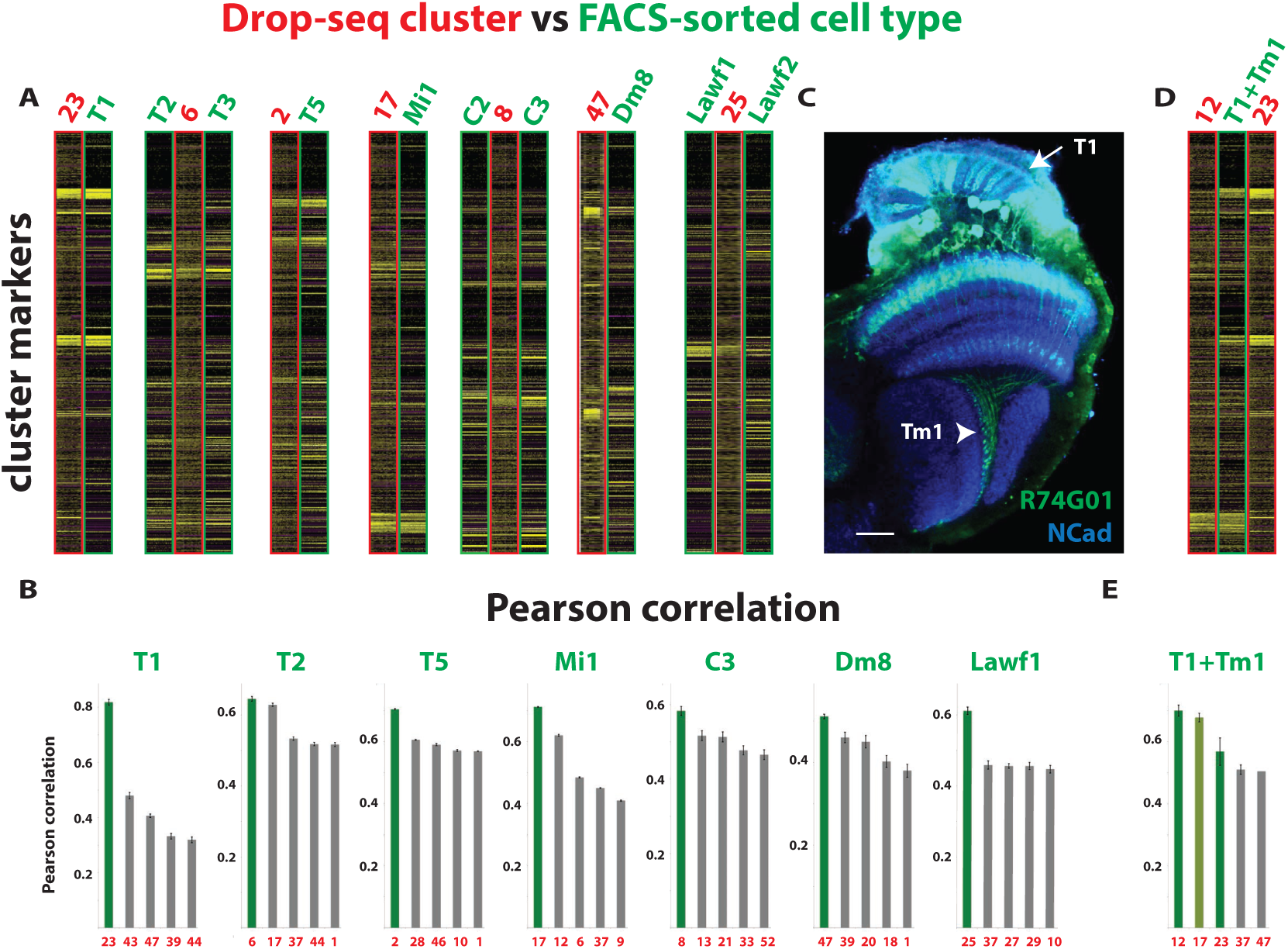
Comparison of Drop-seq cluster transcriptomes and FACS-sorted cell type transcriptomes shows striking similarities between certain clusters and cell types and is used to annotate the Drop-seq clusters. (A) The expression levels of 401 selected Drop-seq cluster markers (rows) is shown for “simulated single cells” (columns) representative of FACS-sorted cell types (green – cell type name indicated on top) and for single cells of the respective Drop-seq clusters (red – cluster number indicated on top). Each FACS-sorted cell type corresponds clearly to one Drop-seq cluster. (B) Histograms showing the Pearson correlation of the transcriptome of each FACS-sorted cell type with the transcriptome of the more correlated clusters. Most of the cell types map to one cluster. Error bars represent standard error of the mean of the triplicates’ Pearson correlation with the more related clusters. (C) *R74G01-Gal4>UAS-myrGFP* is expressed in two cell types: Tm1, whose projections in the lobula are indicated by the arrowhead, and T1, whose projections in the lamina are marked by the arrow. *NCad* is used to visualize the neuropils. Scale bar, 20um. (D) The heatmap shows the expression levels of the 401 selected markers (rows) in “simulated single cells” (columns) that represent R74G01-Gal4 and in single cells of the respective Drop-seq clusters, 12 and 23. Since R74G01-Gal4 is expressed in two different cell types, T1 and Tm1, its transcriptome matches two Drop-seq clusters. (E) Histogram showing the Pearson correlation of the T1 and Tm1 mixed population transcriptome with the more correlated Drop-seq clusters. It maps to two clusters, 12 and 23.

The optic lobe consists of more than 100 different cell types. Since we recovered 52 clusters, some clusters contained cells of more than one cell type. Attesting to the quality of the clustering, we observed that very similar cell types clustered into single clusters, *i.e.* C2/C3 (cluster 8), T2/T3 (cluster 6), T4/T5 (cluster 2), and Lawf1/Lawf2 (cluster 25) (Figure 2A, S2D, and 3B).

**Figure 3:**
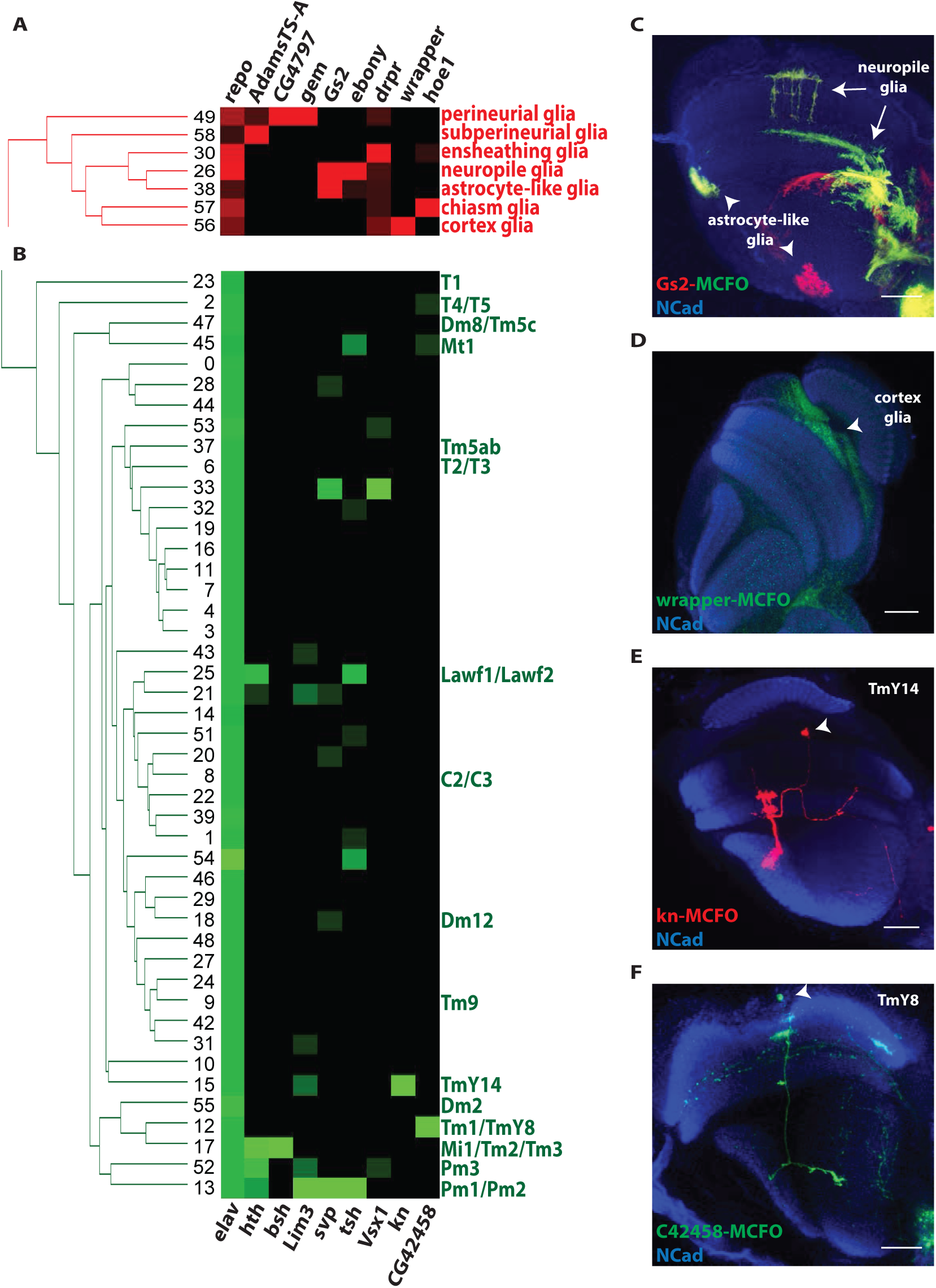
Annotation of glial (red) and neuronal (green) clusters. (A) Annotation of all glial clusters using glial markers. *Repo* is expressed in all glial clusters. *AdamsTS-A* is expressed in perineurial and subperineurial glia, while *CG4797* and *gemini* are only expressed in perineurial glia. *Gs2* is expressed in astrocyte-like glia and neuropile glia (see also Figure 3C). *Drpr* is mainly expressed in phagocytic ensheathing glia. *Wrapper* is only expressed in cortex glia (see also Figure 3D) and *hoe1* mainly in chiasm glia (see also Figure S3C) (B) Annotation of neuronal clusters using three different techniques: 1) based on their correspondence to the FACS-sorted cell type transcriptomes (see also Figure 2), 2) based on known markers (Mi1 expresses *bsh* and Pm1, Pm2, and Pm3 express *Lim3,* Pm1 and Pm2 express *svp,* Pm1 expresses *tsh,* and Pm3 expresses *Vsx1),* 3) based on newly identified markers *(kn* is expressed in cluster 15 and corresponds to TmY14, and *CG42458* is mainly expressed in cluster 12 and corresponds to TmY8). (C) A swapped MIMIC line expressing Gal4 in the pattern of *Gs2* was used to drive MCFO (Nern et al., 2015). Single cell clones were generated in the adult brain and are shown in red and green. *Gs2* is expressed in neuropile glia (arrow) and astrocyte-like glia (arrowhead). (D) A wrapper-Gal4 line was used to drive MCFO (Nern et al., 2015). Single cell clones were generated in the adult brain and are shown in green. *Wrapper* is expressed in cortex glia. (E) A swapped MIMIC line expressing Gal4 in the pattern of *kn* was used to drive MCFO (Nern et al., 2015). Single cell clones were generated in the adult brain and are shown in red. *Kn* is expressed in TmY14. (F) A swapped MIMIC line expressing Gal4 in the pattern of *CG42458* was used to drive MCFO (Nern et al., 2015). Single cell clones were generated in the adult brain and are shown in green. *CG42458* is expressed in TmY8 (arrowhead). *NCad* labels the neuropils in C-F. Scale bar, 20um.

To further demonstrate the accuracy of our clustering and its potential to discriminate between different cell types, we wanted to identify instances where a seemingly homogeneous population actually consisted of more than one cell type. For this purpose, we used a Gal4 line that marks two different cell types - T1 and Tm1 (Figure 2C). We FACS-sorted the cells and obtained the cell type-specific transcriptome, as described earlier. We generated “simulated single cell” transcriptomes, plotted a heatmap with the cluster-specific markers, and compared it to the single cell and cluster heatmap. The “simulated single cells” mapped to two different clusters (12 and 23), which correspond to neuronal types Tm1 and T1 (Figures 2D and 2E).

The mapping of the transcriptomes of these single cell types to unique clusters, as well as the splitting of a heterogeneous cell population into its constituent cell types, indicate the robustness of our clustering technique and the reliability of the clusters.

### Identification and annotation of different glial cell types

One of the drawbacks of Drop-seq is that it does not provide the identity of the sequenced single cells and requires other approaches for their annotation. Matching cell type-specific transcriptomes to unique clusters is a perfect way to annotate the clusters for which FACS-sorted cell transcriptomes were available (Figure 2 and Figure 3B). Although this validates the approach, it does not generate new information about the remaining cell types. We thus aimed to annotate the remaining clusters that should correspond to cells that were not in the collection of FACS-sorted cells.

As a simplified starting point, we focused on annotating glial clusters, which account for ~15% of the total number of clusters (7 out of 52). Glia represent ~10% of the cells in the adult *Drosophila* central nervous system. This number varies extensively in vertebrates and reaches 50% of the human brain (Azevedo et al., 2009; Edwards and Meinertzhagen, 2010; Herculano-Houzel, 2014; Kremer et al., 2017). Several glial cell types have been described in the *Drosophila* optic lobes: surface glia (comprising perineurial and subperineurial glia), cortex/satellite glia, ensheathing glia, neuropile glia, astrocyte-like glia, and chiasm glia (Edwards and Meinertzhagen, 2010; Freeman, 2015; Kremer et al., 2017).

We first identified all glial clusters based on the expression of *repo,* a marker expressed universally and exclusively by glial cells (Figure 3A). To annotate the glial clusters, we used available transcriptomic information and reporters. *AdamTS-A* is more highly expressed in surface glia (perineurial and subperineurial glia) as compared to other glia (DeSalvo et al., 2014) and is involved in their functionality as the *Drosophila* blood-brain barrier. *AdamTS-A* was expressed in clusters 49 and 58. *CG4797* and *gemini* are expressed in perineurial glia that correspond to cluster 49 in which they are also abundant (Figure 3A-Figure S3A). Three clusters (26, 30, and 38) correspond to ensheathing glia: two of them contained *Glutamate synthase 2* (*Gs2*) and *Excitatory aminoacid transporter 1 (Eaat1)* and correspond to neuropile glia and astrocyte-like glia (clusters 26 and 38) (Figure 3A, 3C and Figure S3B). The two clusters were further distinguished by the expression of *ebony*, which is only expressed in neuropile glia (cluster 26) (Richardt et al., 2002). Cluster 30 expresses *Draper* and corresponds to ensheathing glia with phagocytic activity (Doherty et al., 2009). *Draper* is also expressed in cortex glia and is found in higher levels in cluster 56, which was annotated as cortex glia. Furthermore, *wrapper* was only found in cluster 56. We used a wrapper-Gal4 line to show that *wrapper* is indeed only expressed in cortex glia (Figure 3D). Finally, *hoepel1* was most highly found in cluster 57, and *hoepel1* is expressed in chiasm glia (Figure S3C). Therefore, we were able to identify all known glial types in our clusters and assign identity to all seven glial clusters based on specific markers found in each cluster (Figure 3A).

### Molecular markers for the characterization and annotation of neuronal Drop-seq clusters

To annotate more unidentified neuronal clusters, we used known and newly identified cell type-specific markers. Although we have a good understanding of how neuroblasts are temporally and spatially specified to give rise to different neuronal types, we do not know which cell types are born at each temporal window by neuroblasts from a given specific spatial origin. The best-studied temporal window is the *hth* window. Neuroblasts that express *hth* are responsible for generating Mi1, Pm1, Pm2, and Pm3 neurons (Erclik et al., 2017). Mi1 expresses *bsh* and *hth* while Pm1, Pm2, and Pm3 express *Lim3* and *hth.* Moreover, Pm1 and Pm2 express *svp,* Pm1 expresses *tsh,* and Pm3 expresses *Vsx1.* We looked for clusters expressing these combinations of genes and identified cluster 17 as Mi1 (which was also identified by the Drop-seq-FACS comparison - Figure 2A), cluster 52 as Pm3, and cluster 13 as Pm1/Pm2 (Figure 3B). Pm1 and Pm2 are very similar cell types, which explains why they map to the same Drop-seq cluster.

To annotate some of the unidentified clusters, we used Gal4 lines for cluster-specific markers. *kn (collier)* is only expressed in cluster 15. We used Multi-Color Flip-Out (MCFO) (Nern et al., 2015) with a kn-Gal4 line to generate single cell clones in the adult brain to identify the corresponding neuron. The line marked only one cell type, which was identified as TmY14 (Figure 3E). Similarly, *CG42458* was highly expressed in cluster 12 and a *CG42458-Gal4* line was expressed in TmY8 (Figure 3F). *CG42458* was also present in cluster 2, which has already been identified as T4/T5 cells, and in cluster 45, which we annotated as Mt1 neurons, as indicated in Figures S3D-D’.

In conclusion, four different assays were used to assess the quality of our single cell data and the subsequent clustering: *(i)* the Drop-seq clusters matched the transcriptomes of FACS-sorted single cell types; *(ii)* all glial clusters were identified; *(iii)* known markers allowed us to identify clusters, and *(iv)* MCFO with Gal4 lines for unknown markers allowed us to define new expression patterns and to identify further clusters. Altogether, we were able to assign 32 cell types to 23 clusters among the 52. This provides an in depth description of the molecular properties of cell types that were previously only known from their morphology or a few molecular markers.

### Transcription factor-based hierarchical clustering is different from whole transcriptome-based clustering

Single cell transcriptomes are too shallow to recover the expression of every transcription factor in every single cell. Having verified the reliability of the clustering and to study which transcription factors regulate downstream effectors and cell type identity, we treated the data as cluster transcriptomes rather than as single cells. The reads of all single cells that belong to one cluster were merged and normalized (see STAR Methods). We first looked at the expression of transcription factors in the clusters. There are two categories of transcription factors: 72 transcription factors were found at similar levels in most clusters (11 ubiquitous and 61 pan-neuronal), while 598 cell type-specific transcription factors were expressed at significantly higher levels in only one or few clusters (Figure 4A). We also performed weighted gene co-expression network analysis (WGCNA) (Langfelder and Horvath, 2008) and observed that the transcription factors of each co-expression module were mostly expressed in one cluster (Figure 4A’ and S4A) and there was no extensive transcription factor fingerprint overlap between cell types that are closely related. This suggests that apparently similar cell types have different compositions of transcription factors, which we will address in more detail below.

**Figure 4:**
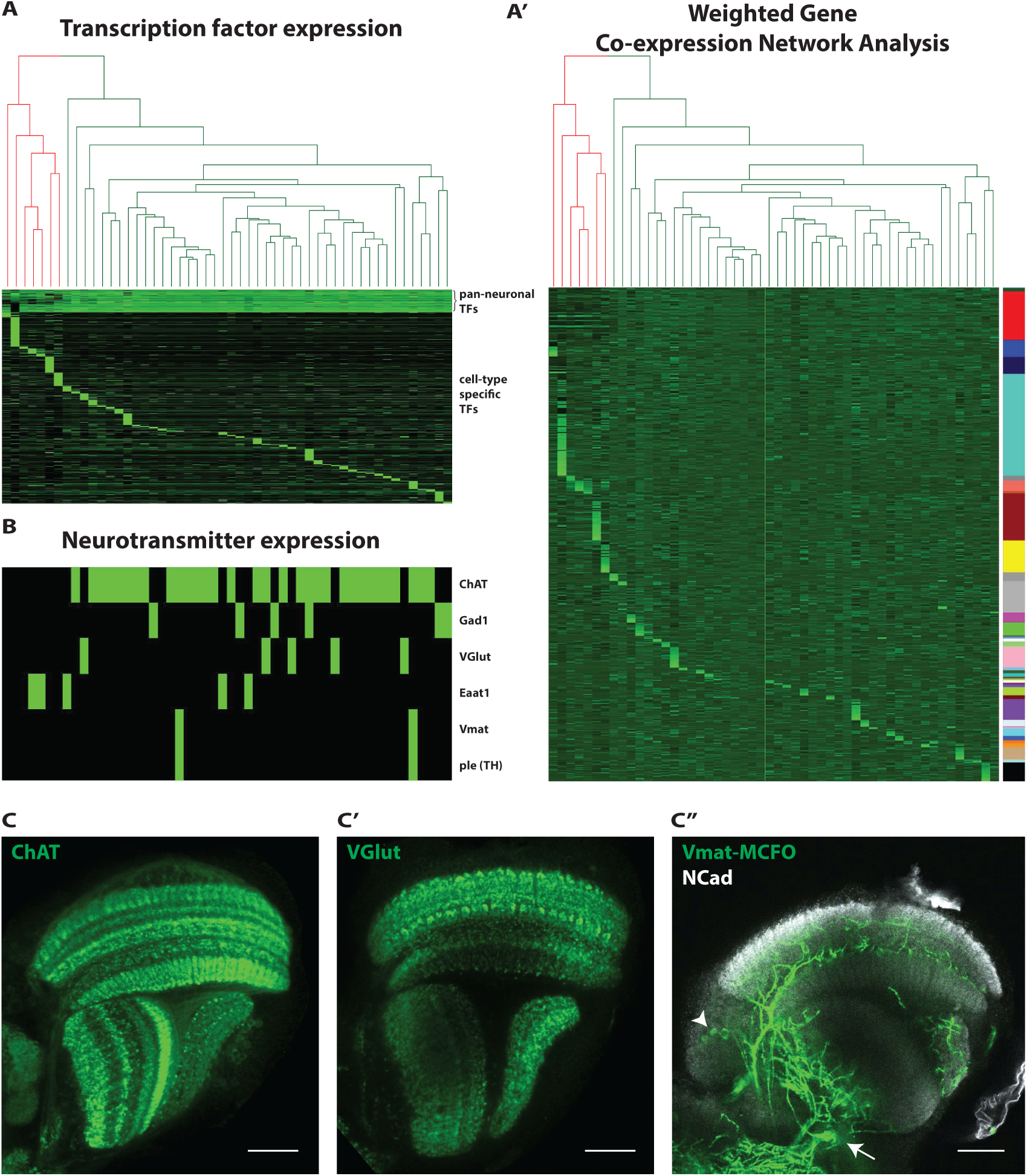
Transcription factor and neurotransmitter expression in the Drop-seq clusters. (A) Heatmap showing the expression of transcription factors (rows) in all Drop-seq clusters (columns) – green in different intensities indicates expression in different levels, black corresponds to no expression. The transcription factors that are expressed in the adult optic lobe neurons and glia can be separated into two categories: 72 ubiquitous/pan-neuronal transcription factors are found at similar levels in most cell types, while 598 cell type-specific transcription factors are expressed in significantly higher levels in only one or few cell types. (A’) Heatmap showing the expression of transcription factors (rows) in all Drop-seq clusters (columns). Transcription factors are organized in modules (color-coded on the right), which were defined by weighted-gene co-expression network analysis (WGCNA). We observe that each module of transcription factors is mainly expressed in a single cluster, indicating that similar cell types have different compositions of transcription factors. (B) Heatmap showing the expression of neurotransmitter related genes (rows) in all Drop-seq clusters (columns) – green indicates expression, black corresponds to no expression. *ChAT* is expressed in cholinergic neurons, *Gad1* in GABAergic, *VGlut* in glutamatergic, *Eaat1* is an excitatory aminoacid transporter that is used to uptake glutamate, *Vmat* marks aminergic neurons and *ple* (tyrosine hydroxylase) is expressed in dopaminergic neurons. Most of the neurons in the optic lobe are cholinergic. (C-C’) Antibody staining against *ChAT* and *VGlut* showing the presence of cholinergic and glutamatergic neurons in the *Drosophila* optic lobe. (C”) A Vmat-Gal4 line was used to drive MCFO (Nern et al., 2015). Single cell clones were generated in the adult brain and are shown in green. Two aminergic neuronal cell types can be seen with their cell bodies in the medulla rim (arrowhead) and the lobula cortex (arrow). Scale bar, 20um.

Having assigned transcription factor fingerprints to each cluster, we sought to determine how they regulate terminal characters. One of the main terminal characters of a neuron is its neurotransmitter identity. We thus analyzed the neurotransmitter composition in each of the clusters. The majority of clusters expressed *Choline acetyltransferase (ChAT)* and was thus cholinergic. Most of the others were either glutamatergic *(Vesicular glutamate transporter* - *VGlut)* or GABAergic *(Glutamic acid decarboxylase 1* - *Gad1)* (Figure 4B). Indeed, antibody staining for *ChAT* (Figure 4C) and *VGlut* (Figure 4C’) showed broad expression patterns in the optic lobes. Two clusters corresponded to neuronal types that appear to be monoaminergic as they expressed the *Vesicular monoamine transporter Vmat* (Figure 4C”) (Nassel and Elekes, 1992). Tyrosine hydroxylase (*ple*) was also expressed in these neurons, indicating that they are likely dopaminergic (Daubner et al., 2011). We did not detect expression of *Tryptophan hydroxylase (Trh)* or *Tyrosine decarboxylase 2 (Tdc2)* in any of the clusters suggesting that serotonergic, octopaminergic, and tyraminergic neurons (Alekseyenko et al., 2010; Cole et al., 2005) do not have cell bodies in the optic lobes. However, some modulatory neurons innervate the medulla as we detected expression of octopamine receptors in different clusters. In fact, there are octopaminergic neurons that reside in the central brain and send their axons to the optic lobe (Busch et al., 2009; Suver et al., 2012).

It is interesting to note that the neuronal cell types did not cluster according to their neurotransmitter profile in the transcription factor-based hierarchical clustering. To address this, we generated a hierarchical clustering tree based on the whole transcriptome rather than only transcription factors: the two trees were different (Figure S4B): The whole transcriptome tree correlated very well with the neurotransmitter composition as glutamatergic and GABAergic neurons now largely clustered with each other (Figure S4B’). This result suggests that although two cell types may have a similar whole transcriptome composition, this similarity is achieved by convergence of different combinations of transcription factors.

### A ‘random forest’ model to predict transcription factors responsible for the regulation of downstream effectors

To directly address the regulation of different genes by transcription factors and establish a causal relationship between transcription factors and downstream targets, we generated a model that could predict a cell type’s gene expression based on its transcription factor profile, and could potentially identify the transcription factors responsible for specific traits.

We randomly chose 39 clusters as a training set and the 13 remaining as a test set (Figure S5A) to train a machine learning ‘random forest’ model to predict the expression level of every gene in the test clusters based on the clusters’ respective transcription factor profile (Figure 5A). The predicted gene expression for the neuronal clusters was 98% accurate with a Pearson correlation ranging from 84.9% to 97.5% between actual and predicted transcriptomes. The performance of the model was lower for glia, with a Pearson correlation of 70.2% (Figure 5A-B). This is probably due to the lower number of glial clusters and, therefore, the incomplete training of the model. To address whether housekeeping genes were mainly driving the high accuracy, we compared the actual and predicted expression of the 1,303 variable genes we had identified earlier (Figure S5B). While the accuracy was still very high for neurons (88%), it was much lower for glia (27%), indicating that more clusters are necessary for training.

**Figure 5:**
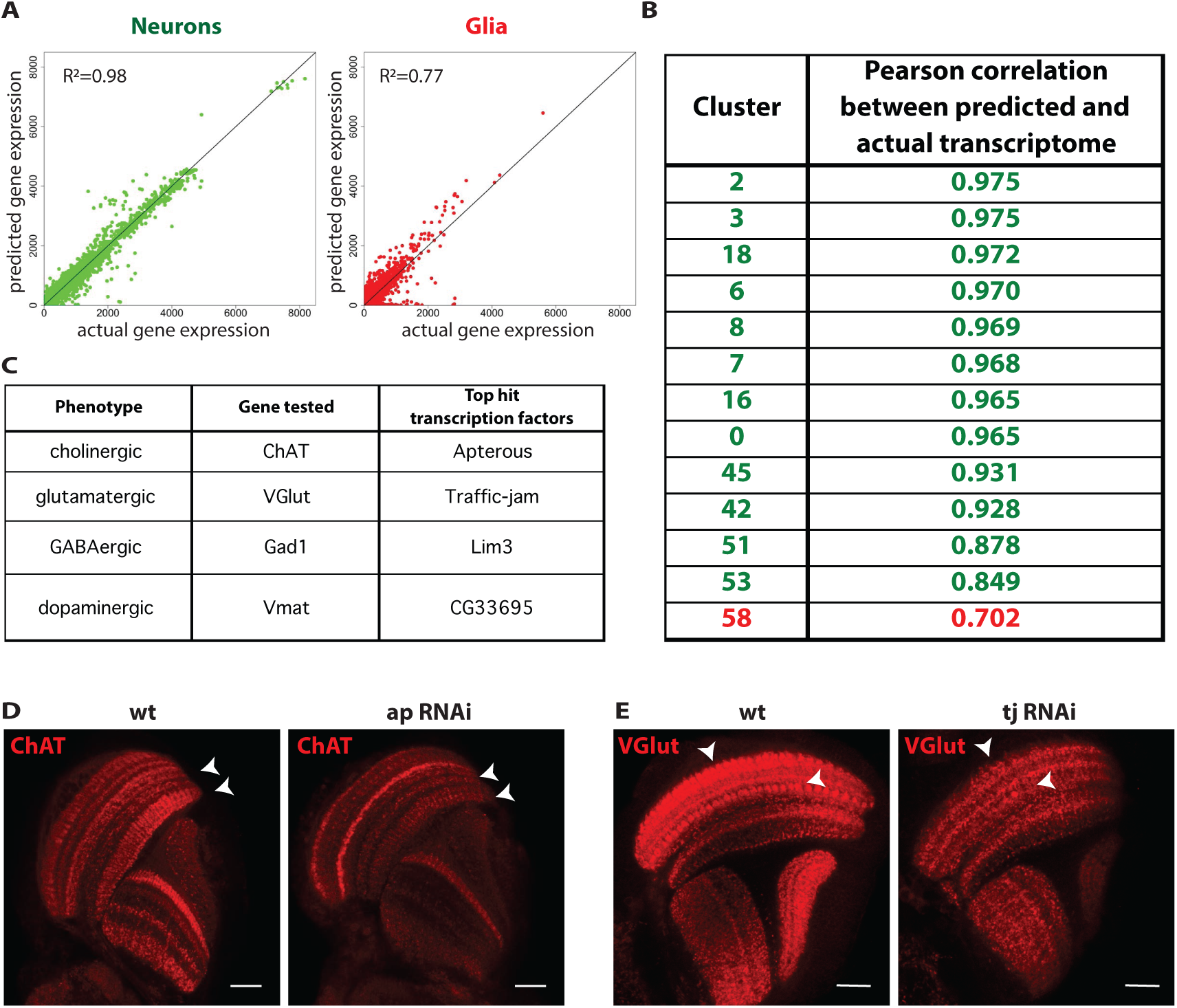
A ‘random forest’ model identifies transcription factors that regulate terminal genes involved in neurotransmitter expression. We trained a ‘random forest’ model using 39 clusters as a training set and 13 clusters as a test set. (A) The generated model can faithfully predict the expression of all genes in the test clusters given the transcription factor expression. The accuracy of the prediction was 98% for the neuronal clusters and 77% for the glial clusters (mainly due to the fewer clusters that led to incomplete training of the model). (B) The Pearson correlation between the predicted and the actual transcriptome ranged from 84.9% to 97.5% in the neuronal clusters and was 70.2% in the glial cluster. (C) Using the ‘random forest’ model, we identified the transcription factors that are mainly responsible for the generation of each of the four neurotransmitter identities: cholinergic *(apterous),* glutamatergic (*traffic-jam*), GABAergic *(Lim3),* and monoaminergic *(CG33695).* (D) The expression of *ChAT* was predicted to rely on the expression of *ap* in a subset of the clusters. Knock-down of *ap* in the adult optic lobe led to the downregulation of *ChAT* in specific medulla layers, M6 and M10, as well as in the lobula. (E) Effect of *tj* knock-down in the expression of *VGlut.* The expression of *VGlut* in synaptic boutons in medulla layers M1 and M6 is reduced upon downregulation of *tj.* Scale bar, 20um.

Using this model, we predicted the transcription factors responsible for generating each of the four neurotransmitters identities found in the optic lobe. The random forest model generated a single score per variable *(i.e.* transcription factor) and we identified the top transcription factors that were predicted to be responsible for cholinergic (*apterous*), glutamatergic (*traffic-jam*), GABAergic *(Lim3),* and monoaminergic *(CG33695)* fate.

To rigorously test the model and verify that we can infer a causal relationship between the identified transcription factors and the neurotransmitter identity, we knocked-down in adult flies the transcription factor that was predicted to regulate a specific gene (to avoid developmental defects or apoptosis during neuronal development) and assessed the effect of the knock-down on the expression levels and pattern of that gene. A heat-shock inducible flip-out actin-Gal4 cassette was activated in adult flies to drive expression of a UAS-RNAi transgene against *apterous.* We performed antibody staining against *ChAT* in the presence and absence of the RNAi. *ChAT* staining was severely reduced in several medulla and lobula layers suggesting that *apterous* is necessary for the expression of *ChAT* in many, although not all, cell types: (Figure 5D). Moreover, we tested the effect of *tj* on *VGlut* expression. Synaptic boutons in medulla layers M1 and M6 that clearly expressed *VGlut* in wild-type optic lobes no longer expressed *VGlut* upon *tj* knock-down (Figure 5E). However, as with *apterous* and *ChAT,* some glutamatergic neurons still expressed *Vglut* after *tj* RNAi expression, suggesting that not all glutamatergic neurons rely on *tj* to regulate *VGlut* expression.

These results show that the ‘random forest’ model can predict transcription factors responsible for regulating downstream effectors. The relationships that we tested were causal, as knocking down the predicted transcription factor eliminated the expression of genes in specific cell types.

### How do transcription factors generate cell types with specific characteristics?

As exemplified by the expression of neurotransmitter genes, we found that a single transcription factor is not necessarily responsible for the expression of a given trait in all cell types (Figures 5D-E). Hence, we set to identify the transcription factors that regulated the expression pattern of given neurotransmitter genes in cell types where the highest-scoring transcription factor is not expressed. *CG16779* and *charlatan* were predicted by the ‘random forest’ model to regulate *ChAT*’s expression pattern in neurons where *apterous* was not expressed. Similarly, the model predicted three other genes for *VGlut* besides *tj (forkhead domain 59A, CG32105,* and *CG4328),* one more gene for *Gad1* besides *Lim3 (eyeless).* We only found one gene for *Vmat (CG33695)* (Figure 6A-A’”).

**Figure 6:**
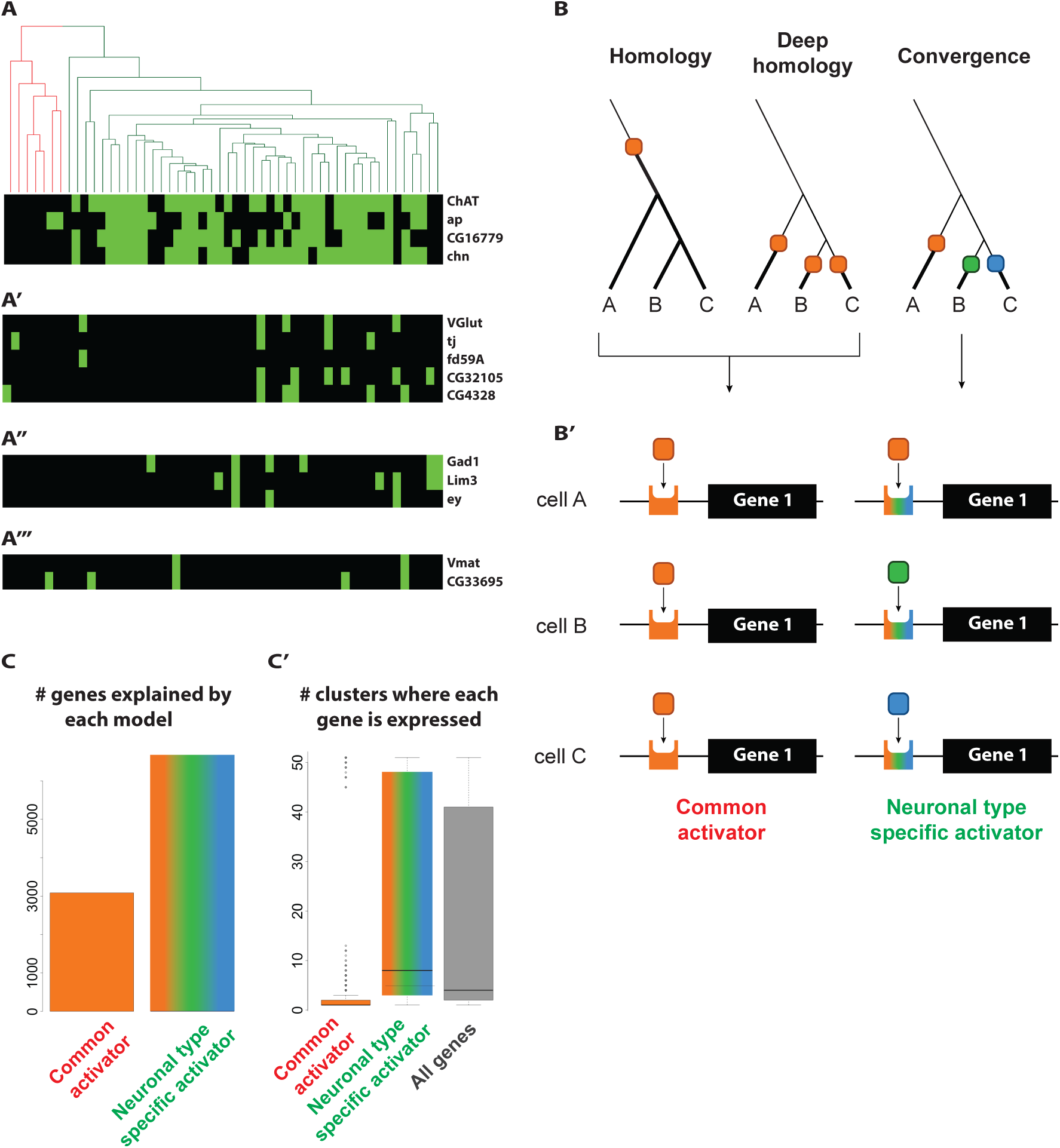
Cell type-specific transcription factors regulate terminal genes in different cells. (A) We used the ‘random forest’ model to identify the top transcription factors that better correlated with the expression of effector genes. *ChAT* expression correlated best with the expression of three transcription factors, *apterous, chn,* and *CG16779.* (A’) Similarly, the expression pattern of *VGlut* may be generated by the combination of four different transcription factors (*traffic-jam, fd59A, CG32105,* and *CG4328).* (A”) The candidates for generating *Gad1* expression pattern are *Lim3* and ey. (A”’) One transcription factor, *CG33695,* was needed to explain the expression of *Vmat.* (B) Two extant cell types that express an effector gene (indicated by the bold line) may either share a common ancestor that was expressing this gene or they may have independently evolved the capacity to express it. In the latter case, the expression of this gene may rely on the same or different transcription factor. (B’) As a consequence of the evolutionary history of the effector gene, its expression in different cell types of the optic lobe may either rely on the same transcription factor (single activator) or different transcription factors may regulate its expression in different cell types (neuronal type-specific activator). (C) Out of the 9738 genes that are expressed in the adult *Drosophila* optic lobe (excluding transcription factors), 3085 genes support a single activator model, while 6653 genes are better explained by a cell type-specific activator model. (C’) The genes that supported the single activator model are expressed in few clusters (2.2 clusters on average), while the 6653 genes explained by the cell type-specific activator model covered a larger range of clusters (22.2 on average).

We wondered whether the regulation of the same gene by different transcription factors in different cell types is a general phenomenon in the fly optic lobes. Two cell types that share common traits (e.g. produce the same neurotransmitter) may either have inherited this trait from a common ancestor, or they may have acquired it independently (Figure 6B). In the first case, the trait would be regulated by the same transcription factor(s) in the two cell types. In the latter case, the trait could be regulated by different (convergence) or by the same transcription factors (deep homology (Tschopp and Tabin, 2017)) (Figure 6B’). We addressed this question by correlating the expression of transcription factors in different clusters with all genes of the same cluster. If a single transcription factor is responsible for the expression of a specific gene, their expression distribution across all clusters should be correlated.

To distinguish between the two models, we selected a gene and asked which transcription factor or combination of transcription factors (among the ~600 that are expressed in the optic lobe) could best recapitulate its expression pattern. We repeated this for every gene expressed in the optic lobe (9738 genes excluding transcription factors), and asked which model (Figure 6B’) was better supported by each of these genes. 3085 genes were better explained by a single activator model, while 6653 genes were better explained by a model where different transcription factors are required to generate its expression pattern (Figure 6C). Since many genes are expressed in one or a few cell types, we wondered whether the 3085 genes of the single activator model corresponded to such genes. Indeed, these genes were expressed on average in 2.2 clusters, while the 6653 genes were expressed in a larger number of clusters (22.2) (Figure 6C’). We also performed Gene Ontology analysis and found that most of the genes related to neural development and differentiation were in the gene pool better explained by the cell type-specific activator model.

These results strongly suggest that most neural traits are regulated independently by different combinations of transcription factors in different cell types. They also corroborate our hypothesis regarding the extensive degree of convergence that explains the difference between transcription factor-based and whole transcriptome-based hierarchical clustering trees.

## Discussion

In this study, we generated a ‘random forest’ model that can successfully predict the expression of a gene based on the cell’s transcription factor profile. In parallel, using this model we identified transcription factors that are responsible for the expression of different effector genes in different cell types. We used RNAi to establish a causal relationship between the expression of the transcription factor and its downstream target. Two of the transcription factors that regulate GABAergic *(Lim3)* and cholinergic (ap) identity are LIM homeobox-containing transcription factors, whose interplay has been shown in both vertebrates and invertebrates to regulate neurotransmitter identity (Hobert and Westphal, 2000; Pfaff et al., 1996; Thor et al., 1999; Thor and Thomas, 1997; Way and Chalfie, 1988; Wenick and Hobert, 2004; Zhang et al., 2014). Interestingly, Lim3 has been associated with either glutamatergic or GABAergic neurons in different systems (Bretzner and Brownstone, 2013; Joshi et al., 2009; Ladewig et al., 2014; Serrano-Saiz et al., 2013; Thor et al., 1999), indicating plasticity, which agrees with our model of extensive phenotypic convergence (Figure 6B). Our data represent an invaluable resource for studying gene regulatory networks and identifying potential terminal selectors and the effectors they regulate.

We do not expect this model to be able to perform well, *i.e.* infer causal relationships, for genes that are expressed in one or very few clusters, because of the lack of proper training. The accumulation of more data from different neuronal systems in *Drosophila,* as well as the constantly improving transcription factor binding site prediction algorithms will help improve the accuracy of our model.

Our study has also provided evidence regarding the mechanisms by which different transcription factors regulate effector genes in diverse cell types. Two main models could be envisaged regarding the utilization of transcription factors for the regulation of effector genes. *(i)* A single (or specific combination of) transcription factor(s) could be regulating the expression of a gene (or gene battery) in every cell type where this is necessary. In this case, every time a gene has to be expressed, this transcription factor is recruited. *(ii)* Distinct transcription factors (or combinations) are used in different cell types to drive the expression of the effector genes. We show that the second model applies to most of the genes required for the function of the *Drosophila* optic lobe neurons, such as neurotransmitters (Figure 6B’). In *C. elegans,* although expression of genes responsible for the generation and release of dopamine in a series of fairly homogeneous dopaminergic neurons expression depends on a single conserved transcription factor *(Ets)* (Flames and Hobert, 2009), the expression of *VGluT* in different neurons appears to rely on multiple regulatory modules in its upstream region and, consequently, distinct transcription factor combinations must control *VGluT* expression in distinct glutamatergic neurons (Serrano-Saiz et al., 2013). Similarly, cholinergic traits are controlled by distinct *C. elegans* transcription factor combinations (Kratsios et al., 2011; Pereira et al., 2015; Wenick and Hobert, 2004; Zhang et al., 2014), and so are GABAergic traits (Gendrel et al., 2016). Here, we present multiple lines of evidence that phenotypic convergence is a more general phenomenon than has been described in worms. The utilization of the same regulatory mode in nematodes and arthropods hints towards a universal strategy for the generation of neuronal diversity that likely also applies to vertebrates.

When two cell types share phenotypic similarity, it is more parsimonious to attribute it to cell type homology and inheritance of the trait from a common ancestor (Figure 6B). In contrast, our results identify significant phenotypic convergence: different cell types use different transcription factors to regulate the very same genes and achieve phenotypic similarity. This is corroborated by the comparison of the hierarchical clustering trees generated using either the whole transcriptome, or transcription factors only. We therefore consider the transcription factor-based tree a better indicator of the developmental or evolutionary history of a neuronal type, although we cannot distinguish between ontogeny and phylogeny (Arendt et al., 2016). In contrast, the whole transcriptome hierarchical clustering is influenced by convergence and delineates functional similarities between adult neurons.

Our data indicate that Drop-seq, despite its shallow transcriptomic information per single cell, can produce meaningful clusters that correspond to single cell types (or, in some cases, a few highly related cell types), when coupled with powerful clustering algorithms. Over-clustering is an important concern when treating single cell data. For this purpose, we complemented single cell sequencing results with traditional RNA sequencing of FACS-sorted cell types to select clustering parameters and to quality control the clustering of the single cells. We further validated the clustering using *Drosophila* genetic tools and pre-existing knowledge.

While exploring the data, we made a number of interesting observations that it will be important to analyze in more detail:

i. Two of our clusters express more than one neurotransmitter - cluster 20 expresses acetylcholine and glutamate, while cluster 54 expresses acetylcholine and GABA. Although there have been indications of neurons co-releasing GABA or glutamate with acetylcholine (Gao et al., 2008; Gendrel et al., 2016; Raghu and Borst, 2011; Ren et al., 2016; Seal and Edwards, 2006; Serrano-Saiz et al., 2017; Tritsch et al., 2016), it is unknown if this is the case in the *Drosophila* optic lobe. An alternative possibility is that these clusters contain more than one cell type that are highly related but differ in their neurotransmitters. Although the number of recovered clusters were sufficient to answer our questions, it will be important for future projects to accumulate more data to divide the clusters that consist of more than one cell type. Resolving clusters 20 and 54 into their eventual constituent cell types will help us address whether there is co-release of neurotransmitters in optic lobe neurons.
ii. Having the transcriptome of every single cell type will give us the opportunity to generate possible neuronal circuits based on the expression of matching cell adhesion molecules in potential synaptic partners. Superimposing these data on connectomic information will allow us to address long-standing questions regarding optic lobe neuronal circuits. Moreover, since synapses are established earlier in development, it will be important to complement our data with cell type-specific transcriptomes at earlier times during pupal development.
iii. There is a neuronal type (cluster 14) whose whole transcriptome resembles glial cells more than neurons (Figure 1E). Like T1, it expresses *Eaat1,* the glial glutamate transporter. Despite its transcription factor identity, the absence of *repo* expression and the expression of *elav* suggest that this cell type is a neuron. It will be interesting to determine its identity and role in the optic lobe. Given that generation of neurons from glia has been reported several times (Bernardos et al., 2007; Doetsch, 2003; Gallina et al., 2016; Sammut et al., 2015), one cannot exclude the presence of such a mixed neuronal type in the adult *Drosophila* optic lobe. Moreover, the astrocyte-like glia cluster shows markedly increased expression of *deadpan* (a neuroblast marker), which agrees with the capacity of glial cells to play the role of neural stem cells.

Finally, we generated cell type-specific transcription factor fingerprints. Given that transcription factors may be considered the drivers of cell type evolution (Achim and Arendt, 2014; Arendt, 2008; Arendt et al., 2016), this can set the framework for future transcriptomic comparative studies that will show the evolutionary history of neuronal diversity in the optic lobe of invertebrates.

## Author Contributions

Conceptualization, N.K, K.K., and C.D.; Methodology, N.K, K.K., R.S., and C.D.; Investigation, N.K, K.K., C.F., L.B.; Writing - Original Draft, N.K, K.K., and C.D.; Writing – Review & Editing, all authors; Supervision, R.S. and C.D.

## Acknowledgements

We are indebted to the fly community for gifts of antibodies and fly stocks, to H. Bellen, Michael Reiser, CH Lee, T.Cook for fly lines, and to H. Aberle for the VGlut antibody. We are very grateful to A. Butler for help with the Seurat package and A. Powers for help with the Drop-seq setup. We also want to thank the NYUAD sequencing and bioinformatics core. Finally, we thank the Desplan and Satija lab members for critical discussion, comments on the manuscript, and support. This work was supported by grants from the NIH (R01 EY017916) and from the NYUAD institute (G-1205C) to CD, and by NIH DP2-HG-009623 (New Innovator award) to RS. NK was supported by postdoctoral EMBO (365-2014) and HFSP (LT000122/2015-L) fellowships. This study is dedicated to the memory of our friend and colleague, Jean-Philippe Grossier.

## Methods

### Drop-seq experimental procedure

Drop-seq was performed as previously described and following the protocol of the McCarroll lab (http://mccarrolllab.com/dropseq/).

*Drosophila* optic lobes were dissected from Canton-S 3-day old females in PBS and were dissociated into single cell suspension by incubating in 2mg/mL collagenase and 2mg/mL dispase in PBS for 1,5 hours at 25°C. The enzymes were then carefully removed and replaced with PBS + 0.1% BSA. The brains are soft but remain intact if pipetted slowly. The brains were pipetted up and down many times (>200-300) until most large chunks of tissue are dissociated. The cells/tissue were kept cold by putting the tubes in ice. The cells were then filtered using 40um cell strainers. The cells were counted immediately before generating the droplets. The concentration of cells loaded on the microfluidic device was 250 cells/ul.

The lysis buffer was slightly modified to contain 0.2mg/ml Proteinase K. The lysis buffer contained: 0.2M Tris pH 7.5, 20mM EDTA, 50mM DTT, 0.2mg/mL Proteinase K, 6% Ficoll PM-400, and 0.2% Sarkosyl.

The droplets (diameter=125um) were then generated as specified in the Drop-seq protocol and collected in a 50ml falcon tube for a run time of 30 minutes. The emulsions were visualized microscopically in a hemocytometer. 5-10% of the droplets contained a bead and we could observe less than 5% bead-occupied droplets with two beads. After droplet formation, the droplets were incubated in a metal bead bath system at 55°C for 10 min. After incubation, the droplets were broken and the samples were processed as described in the Drop-seq protocol.

The samples from each run (three runs in total) were sequenced in 4 lanes using the Rapid Run mode of HiSeq 2500.

### Drop-seq data analysis: filtering, clustering, elimination of over-clustering, and marker selection

#### Filtering

The Drop-seq data were processed with the standard pipeline available from the McCarroll lab (http://mccarrolllab.com/wp-content/uploads/2016/03/Drop-seqAlignmentCookbookv1.2Jan2016.pdf).

For the downstream analyses, we used Seurat (Satija et al., 2015). To select the minimum gene-per-cell cutoff, we arranged the cell barcodes in decreasing order of size and plotted the cumulative fraction of reads. As we did not observe an inflection point, we used the FACS simulated single cell data to decide the cutoff. We used different cut-offs ranging from 100 to 400 genes/cell and assigned each single cell to clusters. We then measured how many of the single cells were correctly assigned. The highest accuracy (~88%) was achieved when selecting cut-offs of 200 and 300 genes/cell. We selected to keep all cells that had more than 200 genes/cell to maintain the greatest number of single cells. We also filtered cells enriched for mitochondrial gene expression (>20%), which is indicative of stressed cells.

#### Clustering

For the clustering, we used a k-nearest neighbor algorithm. To decide on the ‘granularity’ of the clustering (resolution parameter), we used the FACS simulated single cell data again and clustered them using different ‘granularities’. We used resolution values ranging from 0.6 to 8, and we observed that when the resolution was between 4 and 8, we were able to recover unique cluster for every cell type. The accuracy of the clustering was higher when using resolution values 4 and 6. For this purpose, we decided to continue with a resolution value of 4.

#### Elimination of over-clustering

After clustering the single cells, we assessed whether some of the produced clusters were the result of overclustering. We used Seurat to train ‘random forest’ classifiers for each of the terminal nodes and calculated the specificity of the classifier. Nodes with a classifier error higher than 15% were assessed separately for differentially expressed genes, using a likelihood-ratio test for single cell gene expression (McDavid et al., 2013) (Figure S1). In all cases, we were able to recover many highly significantly differentially expressed genes, but we only kept the nodes where the differentially expressed genes were transcription factors, neurotransmitters, cell adhesion molecules, all three of which are indicative of different neuronal types (Figure S1), or well known markers.

#### Marker selection

To select markers for each of the clusters, we used Seurat to perform a ROC test and calculate the ‘classification power’ for each individual marker. We selected the ten markers with the highest classification power for each of the clusters. To compare the Drop-seq single cell data and the FACS-sorted cell type RNA sequencing data, we selected the markers that were expressed in both datasets. We constructed the heatmap of Figure 1 and all downstream applications with this shared marker set.

### Cluster transcriptome analysis

The single cluster transcriptome was generated by adding the counts of all single cells that constituted this cluster and calculating reads per million for each cluster.

For illustrations of Figures 4 and 6, we used binned data. For this purpose, we binned the expression of each gene according to the distribution of its expression in the 52 clusters and scaled it from 0 to 1. For the binning, we generated a histogram (n=40) for each gene’s expression in the 52 clusters and merged the bins that formed “islands” (i.e. they were separated from other “islands” by bins of zero size). We ended up with 2 to 17 bins for each gene that were scaled from 0 to 1 for illustration purposes.

### FACS experimental procedure

We identified lines for individual cell types and crossed each Gal4 line to UAS-Red Stinger to label the nuclei of the specific neuronal type. Dissected brains were dissociated and cells expressing the transgene were sorted on the basis of their fluorescent signal using FACS (FACS Aria III). Cells were sorted directly into extraction buffer and we extracted total RNA using the Arcturus PicoPure RNA Isolation Kit (Applied Biosystems). We assessed RNA quality by Bioanalyzer using RNA 6000 Pico chips (Agilent). Smart-Seq v4 Ultra Low Input RNA Kit was used to generate full-length double stranded cDNA with 300 to 500 pg of total RNA input.

Libraries were prepared using Illumina Nextera XT DNA Library Prep Kit and sequenced on the Illumina HiSeq 2500 System. Three barcoded libraries were pooled per sequencing lane and paired-end 100 bp reads were generated.

Three biological replicates were obtained for each cell type.

### Simulated single cell generation

To compare shallow Drop-seq generated single cell transcriptomes with deep FACS-sorted cell type bulk RNA sequencing, we generated simulated single cells from the FACS-sorted cell types. We generated 900 single cells for each cell type (300 from each of the triplicates). To calculate the number of reads for each simulated cell, we picked a random integer from a normal distribution with the same mean and standard deviation as the number of reads of all Drop-seq generated single cells (i.e. mean = 520.8779, sd = 316.0117). When this number was larger than 200, we sampled this amount of reads randomly from all the genes, with the probability of picking each gene defined by its expression level in this cell type in the bulk RNA sequencing. The simulated single cells were then treated exactly like the Drop-seq sequenced single cells, using Seurat to only keep all cells with reads for more than 200 genes.

### Antibody stainings

*Drosophila optic* lobes were fixed in 4% formaldehyde for 15 minutes at room temperature. After washing, they were incubated for 2 days with primary antibodies at 4°C. After washing the primary antibody, the brains were incubated with the secondary antibodies overnight at 4°C. The secondary antibodies were washed and the brains were mounted in Slowfade and imaged at a confocal microscope (Leica SP5).

### Multi-color flip out clone induction

To generate the multi-color flip out clones, flies carrying the MCFO-1 construct were crossed to ones carrying the specific Gal4 driver. The embryos were raised at 18°C. Once the pupae hatched, the flies were transferred at 37°C for 5 minutes and were again transferred to 18°C. The brains were dissected and stained 2 days after the clone induction.

### WGCNA

Using a weighted-gene co-expression network analysis (WGCNA), we performed a single block network construction and module detection from which we obtained 40 gene-network modules. Given the manageable size of the dataset, we were able to analyze all genes in a single block as opposed to a block-wise network construction method, by setting the maxBlockSize parameter to exceed the number of genes.

The first step for building the weighted network consisted in computing an adjacency matrix reporting the strength of connection between each pair of genes. The adjacency *a_ij_* of two genes i and j with expression levels *x_i_* and *x_j_* was computed using a power adjacency function given by *a_ij_ = cor(x_i_, x_j_*). Upon analysis of network topology for various soft-thresholding powers, we chose *β* to be the lowest power for which approximate scale free topology is attained (i.e.: the power at which the scale free topology fit index curve flattens out upon reaching a high value).

Following that, genes were clustered by average linkage hierarchical clustering, and modules were identified based on topological overlap measure with a minimum module size of 30 genes.

Among the detected modules, we merged modules with highly correlated module eigengenes (a module eigengene can be considered as an average expression profile of the module). For that, we defined a mergeCutHeight value of 0.25, to merge any two modules whose correlation was greater than 0.75.

### RNAi

Flies carrying the hs-flp, actin-flip-out-Gal4, UAS-GFP constructs were crossed to UAS-RNAi lines. The embryos were reared at 18°C. Once the pupae hatched, the flies were transferred at 37°C for 1 hour and 30 minutes to induce the flip-out and activate Gal4 and were then kept at 25°C for one week. The brains were then dissected and stained.

### Antibody generation

A polyclonal antibody against *ChAT* was generated in guinea pigs. The antibody was generated by Genscript (http://www.genscript.com). The epitope used to immunize the guinea pigs were aminoacids 1-480 of the full length protein:

MASNEASTSAAGSGPESAALFSKLRSFSIGSGPNSPQRVVSNLRGFLTHRLSNITPSDTGWKDSILSIPKKWLSTAESVDEFGFPDTLPKVPVPALDETMADYIRALEPITTPAQLERT KELIRQFSAPQGIGARLHQYLLDKREAEDNWAYYYWLNEMYMDIRIPLPINSNPGMVFPPRRFKTVHDVAHFAARLLDGILSHREMLDSGELPLERAASREKNQPLCMAQYYRLLGS CRRPGVKQDSQFLPSRERLNDEDRHVVVICRNQMYCVVLQASDRGKLSESEIASQILYVLSDAPCLPAKPVPVGLLTAEPRSTWARDREMLQEDERNQRNLELIETAQVVLCLDEP LAGNFNARGFTGATPTVHRAGDRDETNMAHEMIHGGGSEYNSGNRWFDKTMQLIICTDGTWGLCYEHSCSEGIAVVQLLEKIYKKIEEHPDEDNGLPQHHLPPPERLEWHVGPQLQLRFAQASKSVDK

### Quantification and statistical analysis

Images were analyzed with FIJI (https://fiji.sc/). All data are expressed as mean ± the standard error of the mean (SEM). ROC tests were used to identify differentially expressed genes between clusters. Pearson correlation on log data was used to test the correlation between FACS-sorted simulated data and Drop-seq clusters, as well as to evaluate the ‘random forest’ model predictions.

### Bioanalyzer

Data was obtained following the standard published protocol: http://www.agilent.com/library/usermanuals/Public/G2938-90014_KitGuideDNA1000Assay_ebook.pdf (Agilent Technologies).

## Supplementary Figures

**Figure S1:**
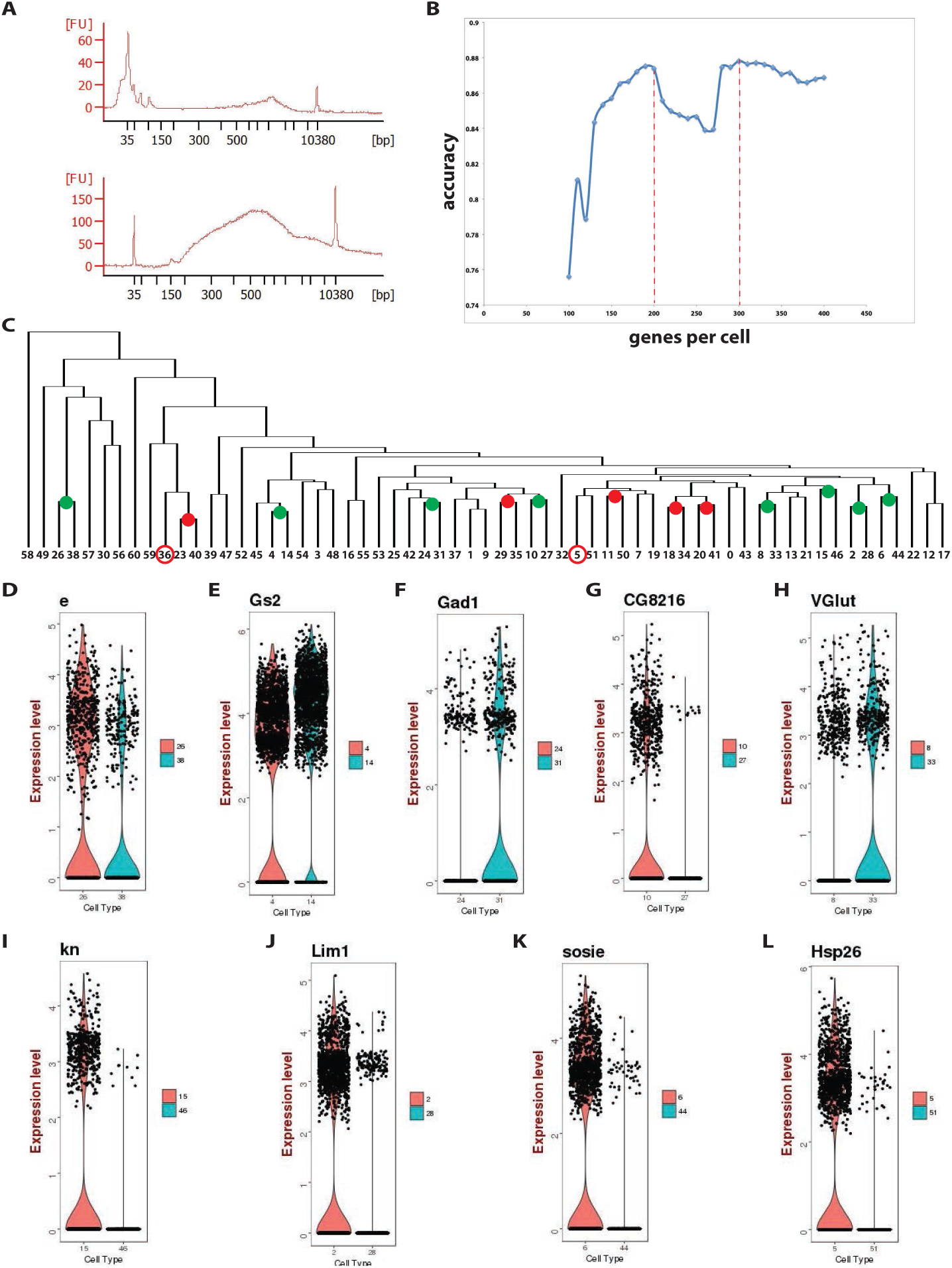
Protocol optimization and parameter selection for the analysis of Drop-seq data. (A) The introduction of Proteinase K in the lysis buffer and a 10-minute incubation of the droplets at 55°C upon droplet formation leads to a >10-fold increase in cDNA yield before library preparation, as shown by the Bioanalyzer electrophoretic trace. FU = fluorescence units, bp = base pairs. (B) To define the minimum amount of genes/cell of each single cell that would be retained for the downstream analysis, we used the simulated single cell data. We selected different cut-offs ranging from 100 to 400 genes/cell and measured the accuracy of assigning the simulated single cells into clusters. The highest accuracy (~88%) is observed with 200 and 300 genes/cell. We selected a cutoff of 200 genes/cell to maintain the greatest number of single cells. (C) Tree of all the clusters using the 1,303 variable genes that were used for the PCA and, therefore, the cluster generation. The validity of each node was assessed by calculating the specificity of a ‘random forest’ classifier trained on each node. The marked nodes indicate an error higher than 15%. The red nodes were collapsed and their clusters were merged (23 and 40, 29 and 35, 11 and 50, 20 and 41, 18 and 34). The encircled clusters were enriched in heat-shock proteins and were therefore eliminated from the downstream analysis. The green nodes were kept based on the differentially expressed genes that were observed between the two clusters (see Figures S1 D-L and STAR Methods). (D-L) Examples of differentially expressed genes between the clusters that were separated by the green nodes. In all nodes, we identified highly significantly differentially expressed genes, but we only kept the nodes where the differentially expressed genes were transcription factors, neurotransmitters, cell adhesion molecules, or well known markers as shown here (see also STAR Methods) (D) *Ebony* was differentially expressed between clusters 26 and 38 (see also Figure 3A), which correspond to two different glial types (astrocyte and neuropile glia, respectively). (E) *Glutamate synthase 2* is differentially expressed between clusters 4 and 14. Cluster 14 is a neuronal cell type that resembles glia in many aspects, as indicated also by the expression of *Gs2,* which is mainly expressed in glia (see also Discussion). (F) *Gad1* is differentially expressed between clusters 24 and 31 (cluster 31 is a GABAergic cell type, while cluster 24 is cholinergic). (G) *CG8216* (space blanket - neuropeptide) is differentially expressed between clusters 10 and 27. Cluster 10 corresponds to cell types Tm5ab. (H) *VGlut* is differentially expressed between clusters 8 and 33 (one is GABAergic, while the other is glutamatergic). Cluster 8 corresponds to cell types C2 and C3. (I) *Knot* is differentially expressed between clusters 15 and 46. *Knot* is a marker gene for TmY14 as shown in Figure 3B. (J) *Lim1* is a transcription factor that is differentially expressed between clusters 2 and 28. Cluster 2 corresponds to cell types T4 and T5. (K) *Sosie* is differentially expressed between clusters 6 and 44. Cluster 6 corresponds to cell types T2 and T3. (L) Heat-shock proteins were enriched in clusters 5 and 36, which were eliminated from downstream analyses.

**Figure S2:**
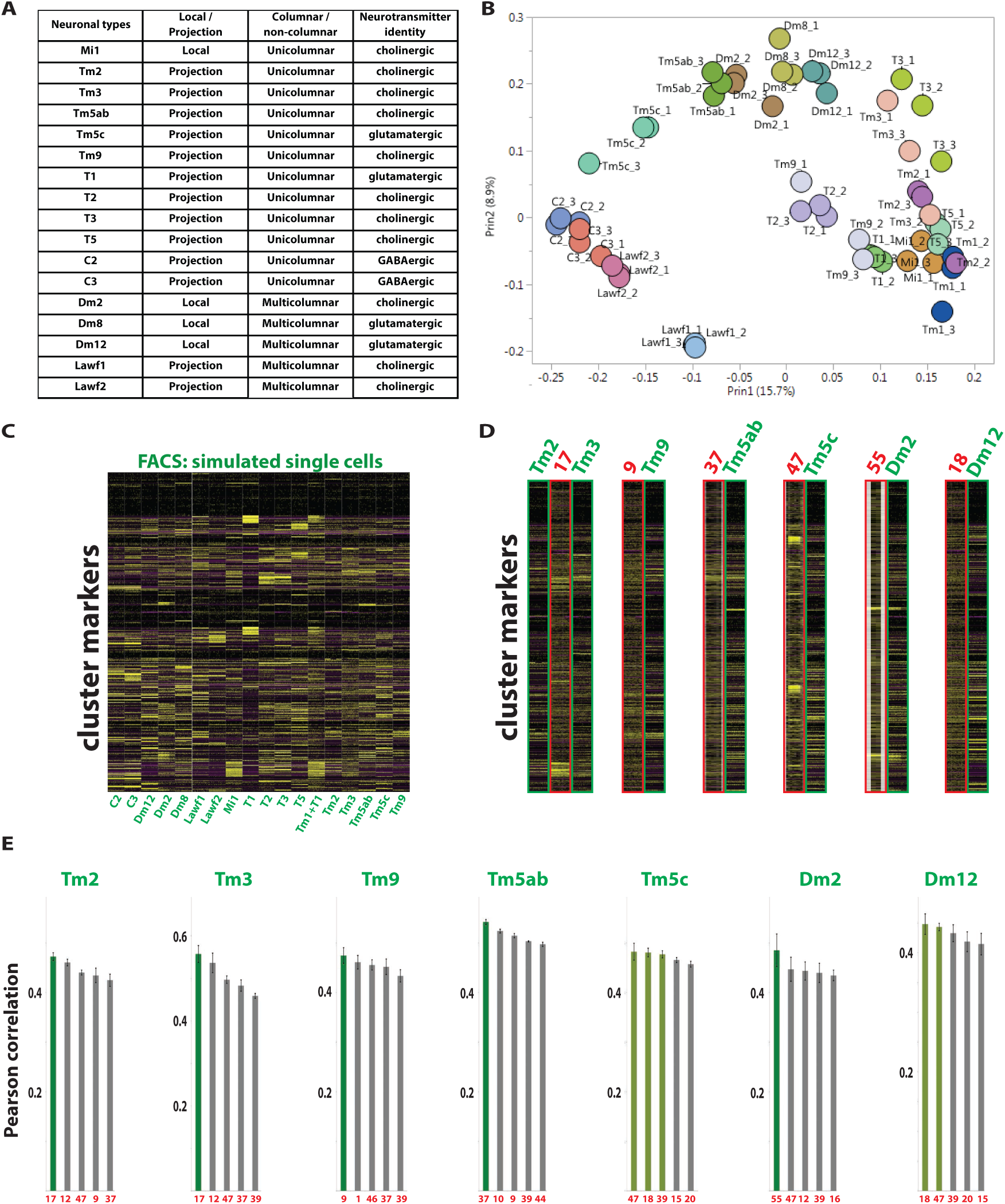
Generation of cell type-specific RNA sequencing data using FACS and comparison to Drop-seq clusters. (A) We used bulk RNA sequencing after FACS sorting to generate the transcriptome of 17 cell types. 4 were locally arborizing, while 13 were projection neurons. 12 were unicolumnar and 5 multicolumnar. 11 were cholinergic, 4 were glutamatergic, and 2 were GABAergic. (B) PCA plot of the biological triplicates of each cell type. (C) The expression of 401 selected Drop-seq cluster markers (rows) is shown in all simulated single cells (columns). Cell types are separated by white lines. (D) The expression of 401 selected Drop-seq cluster markers (rows) is shown in “simulated single cells” (columns) representative of FACS-sorted cell types (green – cell type name indicated on top) and in single cells of the respective Drop-seq clusters (red – cluster number indicated on top). Each FACS-sorted cell type can be assigned to one Drop-seq cluster. (E) Histograms showing the Pearson correlation of each FACS-sorted cell type with the more correlated clusters. Most of the cell types map to one cluster. Tm5c and Dm12 may be assigned to more than one cluster. Error bars represent standard error of the mean of the triplicates’ Pearson correlation with the more related clusters.

**Figure S3:**
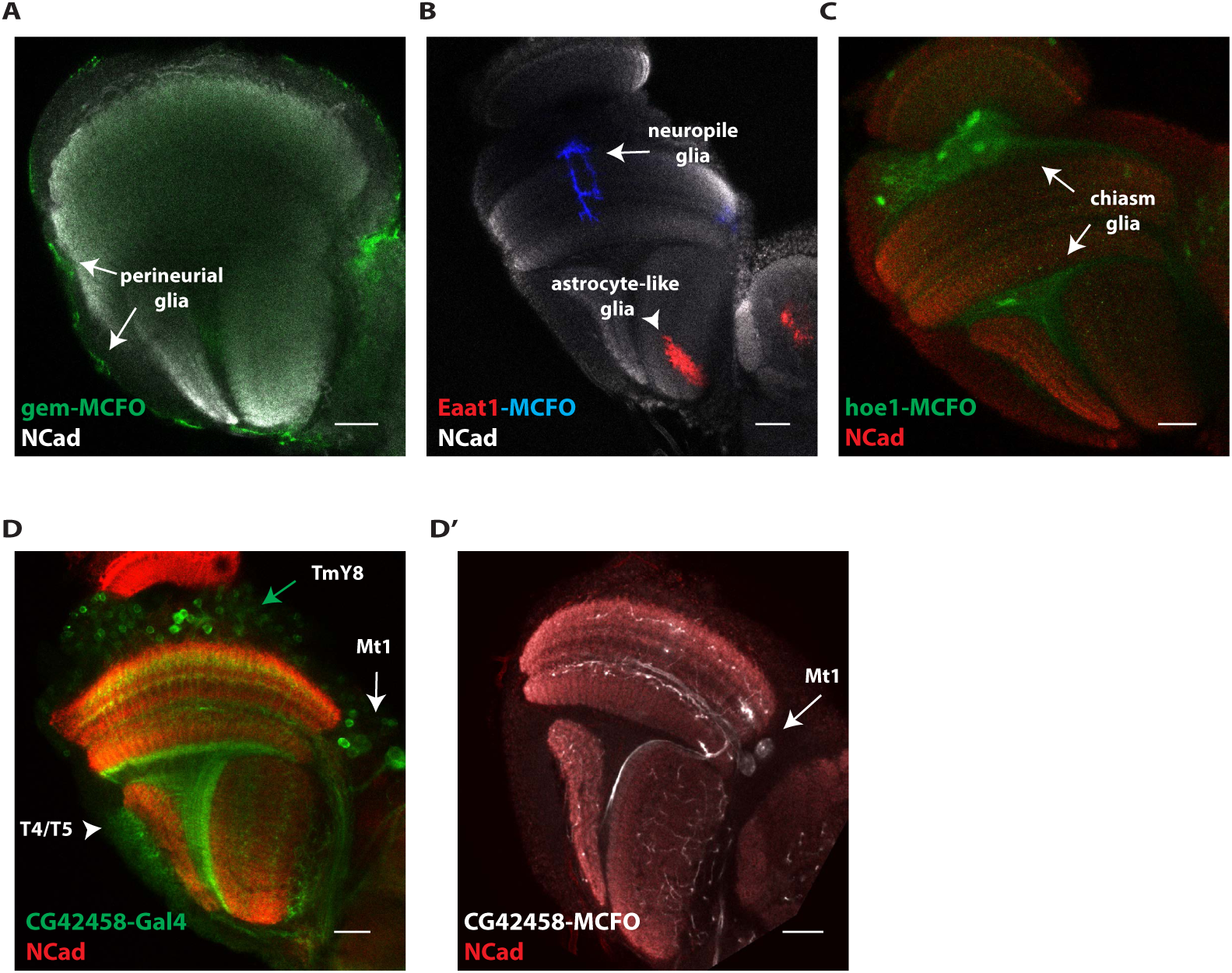
Expression of specific genes in glial and neuronal subtypes was used for cluster annotation. (A) A swapped MIMIC line expressing Gal4 in the pattern of *gemini* was used to drive MCFO (Nern et al., 2015). Single cell clones were generated in the adult brain and and are shown in green. *Gemini* is expressed in perineurial glia (arrows). (B) An *Eaat1-Gal4* line was used to drive MCFO (Nern et al., 2015). Single cell clones were generated in the adult brain and are shown in red and blue. *Eaat1* is expressed in neuropile glia (blue - arrow) and astrocytes (red - arrowhead). (C) A swapped MIMIC line expressing Gal4 in the pattern of *hoe1* was used to drive MCFO (Nern et al., 2015). Single cell clones were generated in the adult brain and and are shown in green. *Hoe1* is expressed in inner and outer chiasm glia (arrows). (D) A swapped MIMIC line expressing Gal4 in the pattern of *CG42458* was used to drive GFP. *CG42458* is expressed in T4/T5 (arrowhead), Mt1 (white arrow), and TmY8 (green arrow). (D’) A swapped MIMIC line expressing Gal4 in the pattern of *CG42458* was used to drive MCFO (Nern et al., 2015). Single cell clones were generated in the adult brain and are shown in white. *CG42458* is expressed in Mt1 cells (arrow). *NCad* labels the neuropils. Scale bar, 20um.

**Figure S4:**
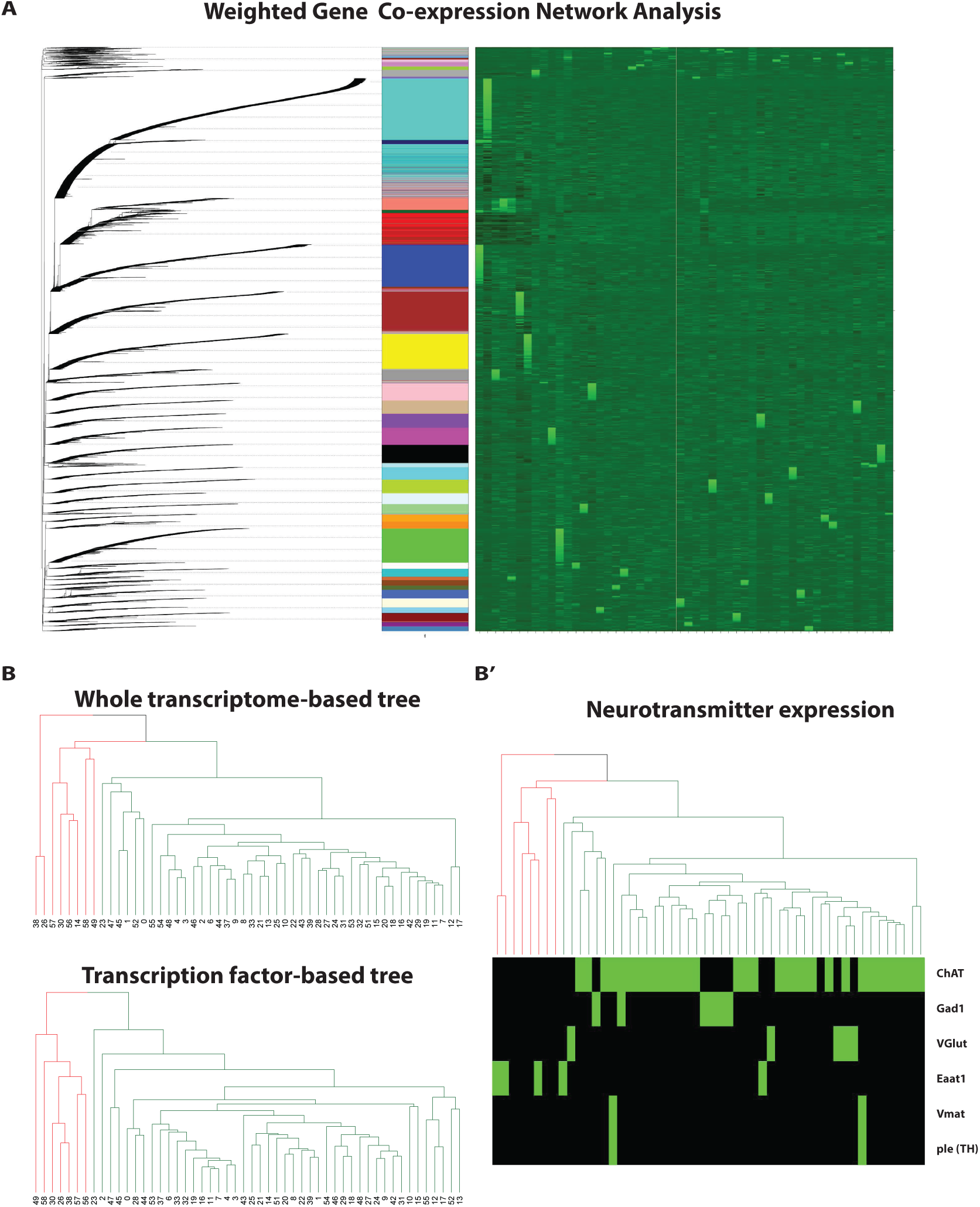
Transcription factor and neurotransmitter expression in clusters. (A) Dendrogram and heatmap of the expression of all genes (rows), organized in 40 modules by weighted-gene co-expression network analysis (WGCNA), across all 52 clusters (columns). (B) Side-by-side comparison of the two hierarchical clustering trees. One was made using the entire transcriptome, while the other one only with the transcription factors. Glia and neurons split early from each other in both trees (in the whole transcriptome tree, neuropile and astrocyte-like glia root the tree). However, cluster 14 clusters with glia in the whole transcriptome tree (although it does not express *repo* – see Discussion). The neuronal cluster distribution in the two trees is very different. (B’) Heatmap showing the expression of neurotransmitter related genes (rows) in all Drop-seq clusters (columns) – green indicates expression, black corresponds to no expression. The tree was generated by hierarchical clustering of the clusters’ entire transcriptomes. In this case, the tree correlates very well with the neurotransmitter composition as most glutamatergic and GABAergic neurons now largely cluster with each other.

**Figure S5:**
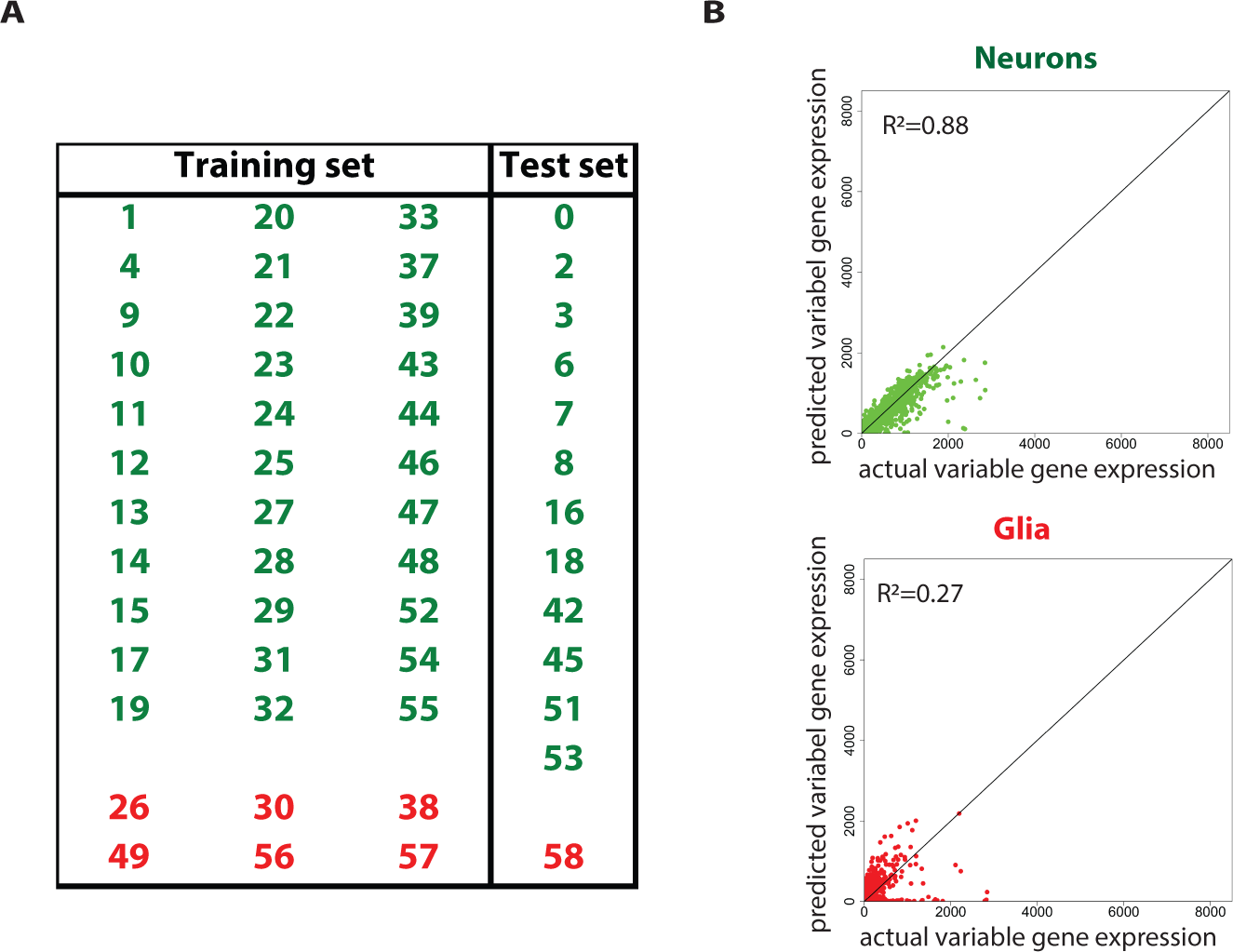
Neuronal and glial training and test sets that were used to test the ‘random forest’ model for the prediction of gene expression based on transcription factors. (A) 33 neuronal clusters (green) and 6 glial clusters (red) were used for training the ‘random forest’ model. The rest were used as a test set. (B) The generated model can faithfully predict the expression of variable genes in the neuronal clusters given the transcription factor expression. The accuracy of the prediction was 88%. The model does not perform well in predicting variable gene expression in glial clusters, probably due to the incomplete training of the model.

